# Distinct competitive impacts of palatability of taste stimuli on sampling dynamics during a preference test

**DOI:** 10.1101/2021.10.10.463786

**Authors:** Benjamin Ballintyn, John Ksander, Donald Katz, Paul Miller

## Abstract

Food or taste preference tests are analogous to naturalistic decisions in which the animal selects which stimuli to sample and for how long to sample them. The data acquired in such tests, the relative amounts of the alternative stimuli that are sampled and consumed, indicate the preference for each. While such preferences are typically recorded as a single quantity, an analysis of the ongoing sampling dynamics producing the preference can reveal otherwise hidden aspects of the decision-making process that depend on its underlying neural circuit mechanisms. Here, we perform a dynamic analysis of two factors that give rise to preferences in a two-alternative task, namely the distribution of durations of sampling bouts of each stimulus and the likelihood of returning to the same stimulus or switching to the alternative—i. e., the transition probability—following each bout. The results of our analysis support a specific computational model of decision-making whereby an exponential distribution of bout durations has a mean that is positively correlated with the palatability of that stimulus, but also negatively correlated with the palatability of the alternative. This impact of the alternative stimulus on the distribution of bout durations decays over a timescale of tens of seconds, even though the memory of the alternative stimulus lasts far longer—long enough to impact the transition probabilities upon ending bouts. Together, our findings support a state-transition model for bout durations and suggest a separate memory mechanism for stimulus-selection.

## Introduction

The ability to efficiently forage for food and other resources is critical for survival, and evolution has therefore doubtless shaped the neural circuitry responsible for decision-making in order to optimize performance of this task (Hayden et al., 2011; Pearson et al., 2014). A key question faced during a foraging bout is a value-based decision, in which the animal uses available information to determine whether to continue to gather food at a current location, or to switch to a potentially more profitable source of nourishment.

Studies of value-based decision-making have typically fallen into two categories: 1) those using “self-control” or “delay-discounting” tasks, in which animals navigate a tradeoff between waiting times and payoff sizes (Bateson & Kacelnik, 1996; Blanchard et al., 2013; Pearson et al., 2010; Stephens, 2002; Stephens & Anderson, 2001); and 2) those using “stay-switch” or “patch-leaving” tasks, in which animals are presented with a source of reward and must decide when to leave it in search of a better alternative (Barack et al., 2017; Blanchard & Hayden, 2014; Constantino & Daw, 2015; Hayden et al., 2011). Patch-leaving tasks better represent the foraging situation (in which animals have sequential interactions with individual reward sources) than do tasks involving the simultaneous presentation of (cued) alternatives, but the ecological realism of patch-leaving studies is limited by the inclusion of a trial-based structure with randomly presented patches; in the wild, in contrast, animals use their experience with the environment to direct their encounters with patches. In some ways, simple two- or multi-bottle preference tests better represent a (simplified) naturalistic foraging scenario, in that an animal can rapidly learn about the state of its environment and direct its encounters toward the alternatives that may have different levels of reward.

Such preference tests are widely used to measure the relative hedonic values—how pleasurable, rewarding, or palatable a taste is, compared to other tastes—of a set of stimuli, with the degree of preference measured in terms of either of the total amounts consumed or the number of licks at a food source. The total amount of a stimulus consumed is a function of: 1) the number of times that stimulus is visited (as compared to the alternative); and 2) the mean amount consumed per visit (or mean duration of visits). But each of these two factors can in turn depend on multiple stimulus properties, including the value of both the visited stimulus and that of alternative stimuli. Thus, a deeper assessment of this type of decision-making requires quantification of these underlying behavioral factors and stimulus properties. One of the goals of this work is to perform this quantification, and to thereby test predictions of a model that explicitly considers such variables (Ksander et al., 2021).

Several questions can be asked about the sequences of decisions that animals make in the process of performing preference tests. Perhaps the most obvious and important is: Do animals rapidly settle on a favorite option, or do they continue to switch between and sample both options even after establishing a favorite? If the animal has a clear preference between two options, then an intuitive and theoretically optimal strategy is to first sample both options, determine a favorite quickly, and then spend all of the remaining time sampling the favored option (or until the source is exhausted or the animal is sated). In such a case, it would be very difficult to even quantify the palatability of a tastant by this method: only a ranking would be possible.

If this is not the policy adopted by the animal (as our data, in accordance with many prior studies, shows it is not), then several additional questions can be asked. For instance, if animals continue to switch back and forth between the two options, it is important to ask how sampling times at one option depend on that option’s palatability—as measured by total amount consumed in sessions without alternatives—and on the palatability of the alternative. To answer this question, one could analyze durations of rhythmic bouts of licking, which are comprised of series of rapid licks without significant pauses, to assess whether and how the behavior at one lick spout depends on the contents of the alternative lick-spout. And if the memory of an alternative available tastant significantly impacts licking, we can probe the timescale of such memory by assessing whether this impact decays over time passed since the alternative was last sampled (such that the animal eventually returns).

Because licking is a highly regular, rhythmic behavior, bouts are easily demarcated by pauses, following which the animal either returns to a new bout of licking at the same spout or switches to the alternative spout. For any pair of stimuli, we can produce a transition matrix, indicating the animal’s likelihood of switching to the alternative or staying with the same stimulus after a pause. Analysis of such transition matrices for pairs of stimuli can reveal how the animal weighs the relative value of each, thereby allowing us to test whether the choice of which stimulus to approach (determined by the transition matrix) and the choice of how long to sample that stimulus (determined by bout durations) reflect the same or different underlying cognitive processes.

It is worth noting that competition can arise in preference tests in the absence of any direct interaction between the hedonic value of one alternative and the behavior displayed at the other. The sources of implicit competition include the limited time available in most tasks and limits in the total amount of food desired prior to satiety. Because of these limits, time spent consuming one stimulus necessarily translates into less time at the alternative, regardless of the transition matrix and bout durations for the alternative. Indeed, one can hypothesize a situation in which an association with a more appetizing stimulus might boost the perceived palatability of a paired neutral stimulus leading to longer bouts at the neutral alternative, even as total amount consumed at the neutral alternative goes down due to the fewer visits there. In contrast, if behavior in preference tests resembles that during foraging, one would anticipate that the greater the value of one stimulus, the less time spent during each visit at an alternative source. A primary goal of this work is to identify the nature of the across-stimulus interaction.

Here, we analyze the behavior of rats engaged in a naturalistic continuous-time taste preference task. We also compare the time course of rat behavior with the dynamics of a simulated circuit of model spiking neurons designed to possess two states, one representing the ongoing choice to sample a stimulus, the other to leave that stimulus. Competition between the bout durations of successive stimuli can arise in the model from adaptation-like processes, which drive a negative correlation between one bout of sampling a stimulus and the subsequent bout with the alternative stimulus. We assess our behavioral findings for evidence of such a competitive interaction.

## Results

### Measurement of palatability

To study the dynamics of stay-switch decision making, rats completed two weeks of sessions in the preference testing chamber (Figure 1A/B). On each day, they were given one hour to freely sample from two delivery spouts a random pair of two solutions, each drawn (with replacement) from a possible four solutions, and each with a distinct palatability (distilled water, dH_2_O; and three concentrations of sodium chloride, NaCl, 0.01M, 0.1M, and 1M) (Sadacca et al., 2012). Licks at each delivery spout were recorded using a custom circuit and identified using a semiautomated process (see Methods).

**Figure 1.**
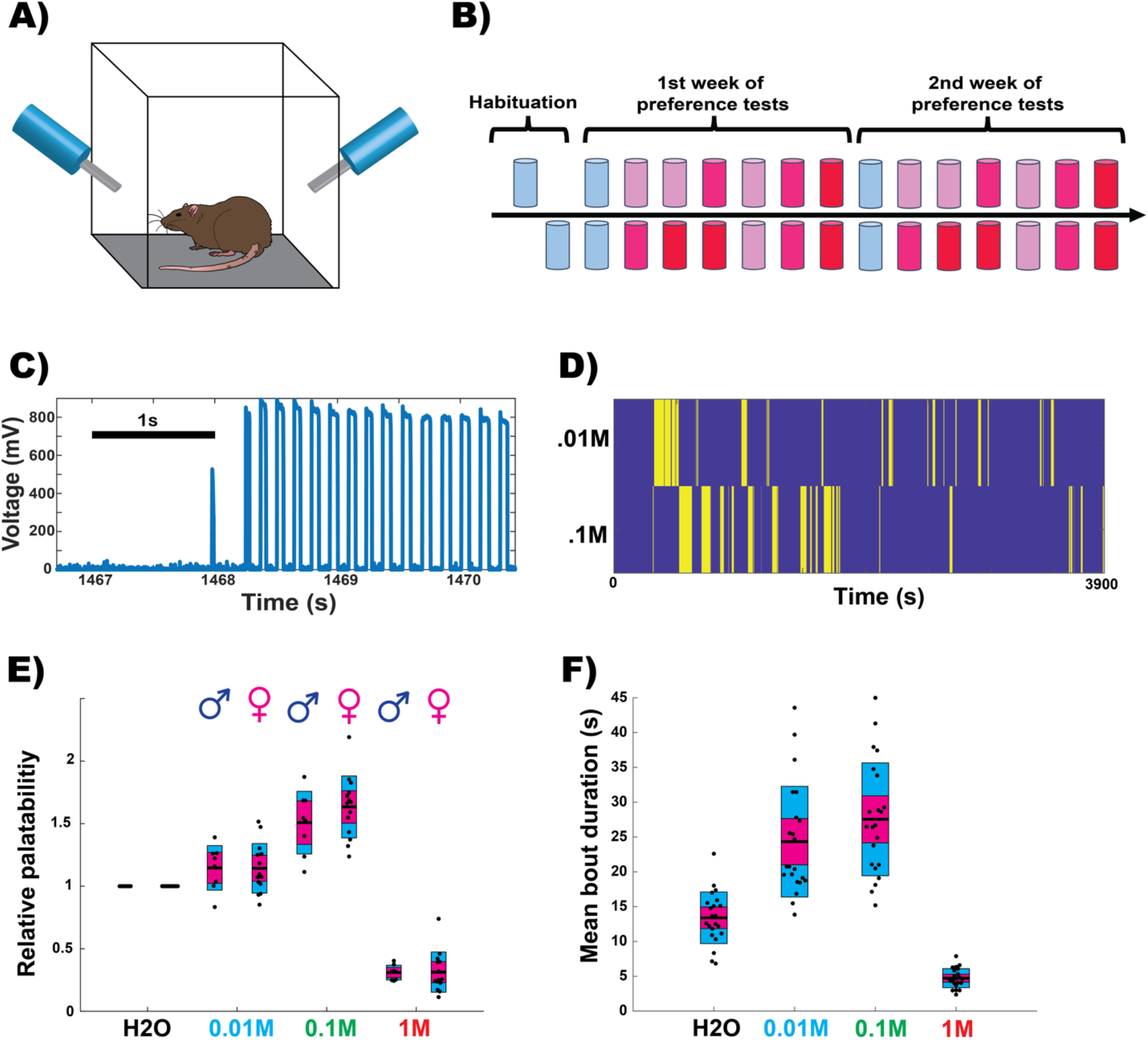
Rats sample fluids in bouts of rapid licking whose durations vary with palatability of the taste stimulus. **A)** 1’ x 1’ custom acrylic chamber had a solution spout available through the left and right walls each containing 25mL of 1 of 3 different NaCl solutions (0.01M, 0.1M, 1M) or dH2O. Rats were allowed to freely move and sample from either spout over the course of 1 hour. **B)** Preference test timeline. Rats were given 2 habituation days with 1 bottle of dH2O on opposite sides across sessions. This was followed by 2 weeks of sessions where each week started with a session of dH2O only followed by all 6 combinations of NaCl solutions. **C)** Example licking data. Each rectangular deflection is one lick. **D)** Example sampling data from a session with 0.01M and 0.1M NaCl solutions. Yellow stripes represent active sampling at the corresponding solution. **E)** Relative palatabilities of the 3 NaCl solutions relative to water. Palatabilities are based on the total number of licks at each solution during sessions where the solution was paired with itself. No sex specific differences were found (0.01M: two-tailed: z = .17, p = .86, .1M: z = −1.06, p = .29, 1M: z = .51, p = .61). Based on pairwise comparisons (the rank order of palatabilities from highest to lowest is (0.1M, 0.01M, H2O, 1M). **F)** Mean bout durations for the 3 NaCl solutions during sessions where each solution was paired with itself. Each datapoint represents the mean bout duration for 1 rat at the corresponding solution.

As our baseline measure of preference, we first confirmed the rank order of the relative palatabilities of the saline solutions (Figure 1E), by measuring the total number of licks to each on days when the solution was paired with itself and dividing by the mean number of licks to dH_2_O on dH_2_O only days. The previously determined palatability ranking (0.1M > 0.01M > 1M; Sadacca et al., 2012) was recapitulated, and as no sex-specific differences were found (two-sample t-test: 0.01M: *t_20_* = 0.042, p = .97; 0.1M: *t_20_* = −1.14, p = .27; 1M: *t_20_* = −0.059, p = .95), data from both sexes were combined for all analyses in which different solutions were pitted against one another (see Methods and Figure 1).

Given the observed differences in palatabilities, we expected that we would also observe different distributions of sampling durations, with more palatable solutions having on average longer durations of lick bouts. This expectation was borne out: the bout duration distributions are mostly commensurate with the calculated palatability of each solution (dH_2_O: mean ±SE = 13.39 ±.79s, .01M: mean ± SE = 24.33 ± 1.69s, .1M: mean ± SE = 27.54 ± 1.72s, 1M: mean ± SE = 4.72 ± .29s). Additionally, the distributions of bout durations of all solutions were well approximated by exponential distributions (Supplementary Figure 1), with decay constants akin to the mean time of bouts at each solution. These exponential distributions of durations are in accordance with the noise-induced state transition model of our computational simulations (Ksander et al., 2021).

### Impact of relative palatability on bout duration

While it is unsurprising that a higher palatability of the currently sampled solution translates into longer sampling bouts, a question that remains unanswered is how the palatability of the alternative solution in a preference test impacts these sampling bout durations. We considered three possibilities: 1) a high alternative palatability will have an appetitive effect, increasing the perceived palatability of the current solution and leading to longer sampling bouts; 2) conversely, a higher alternative palatability could reduce the perceived palatability of the current solution, leading to shorter sampling bouts; and 3) the palatabilities of alternative choices could have no impact on bouts at the current solution. Implied in both hypotheses 1 and 2 is the maintenance of a memory of the alternative solution’s value (palatability) from one bout to the next.

To evaluate the above possibilities, we performed multilinear regression, predicting bout duration as a function of the palatability of available alternatives (mean ± SE *R^2^* = 0.8 ± 0.033, *N* = 22) such that the resulting regression coefficients reveal the impact of the currently sampled and alternative solutions’ palatabilities on the current bout duration. As suggested by differences in mean bout durations across solutions, regression coefficients for the current solution’s palatability were significantly positive (z = 4.09, p = 2.15e-5, mean ± SE = 15.78 ±1.63)—that is, the more palatable a stimulus is, the longer the bouts of licking at it are (Figure 2B).

**Figure 2.**
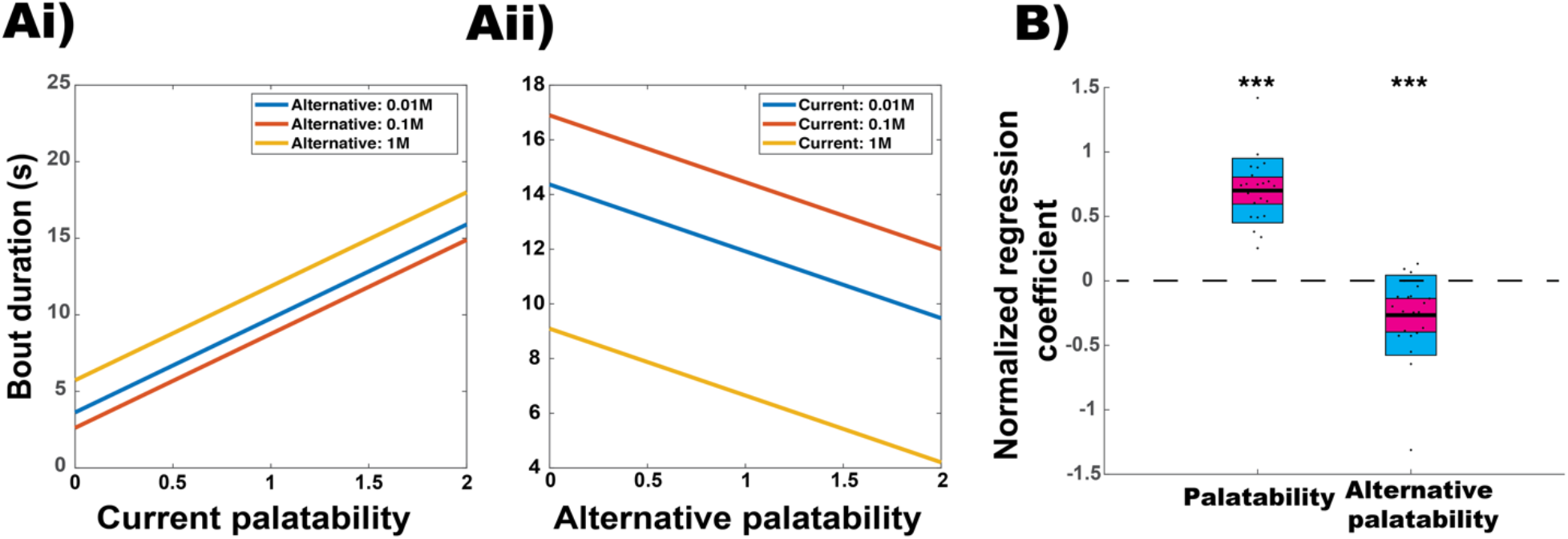
The palatability of the current and alternative stimulus both impact the duration of lick bouts. **A)** Example results from 1 rat of a multilinear regression model for predicting bout duration with the currently sampled solution’s and alternative solution’s palatabilities as factors. **Ai)** Using best fit regression coefficients for 1 rat, bout duration is plotted against the current solution’s palatability for all 3 possible alternative solutions. **Aii)** Same as (Ai), but plotting bout duration vs. the alternative solution’s palatability for 2 levels of the current solution’s palatability. **B)** Current and alternative palatability regression coefficients normalized by the mean bout duration for each rat. Normalized coefficients for current palatability are significantly positive (right-tailed: z = 4.09, p = 2.15e-5) and those for alternative palatability are significantly negative (left-tailed: z = −3.57, p = 1.7e-4).

Alternative palatability coefficients were found to be significantly negative (mean ± SE = −5.69 ±1.32), consistent with possibility 2 above (Figure 2B)—durations of bouts are shorter when the alternative stimulus is of higher palatability.

These results were stable across the course of the session, despite eventual reductions of bout durations due to satiation. When we split sessions into ‘early’ and ‘late’ portions based on a per-animal criterion (we used the 2^nd^ derivatives of each rat’s cumulative distribution of lick times to detect the “kink” in the curve of Figure 3A, where licking slowed from a high rate to a lower rate) and performed the same multilinear regression on early/late bouts separately (early: mean ± SE *R^2^* = 0.66 ± 0.047, late: mean ± SE *R^2^* = 0.77 ± 0.045), we found no significant change between the early and late portions of the session in regression coefficients of normalized bout durations (Figure 3B/C). Higher palatability of the available alternative translates into shorter licking bouts at the spout in front of a rat.

**Figure 3.**
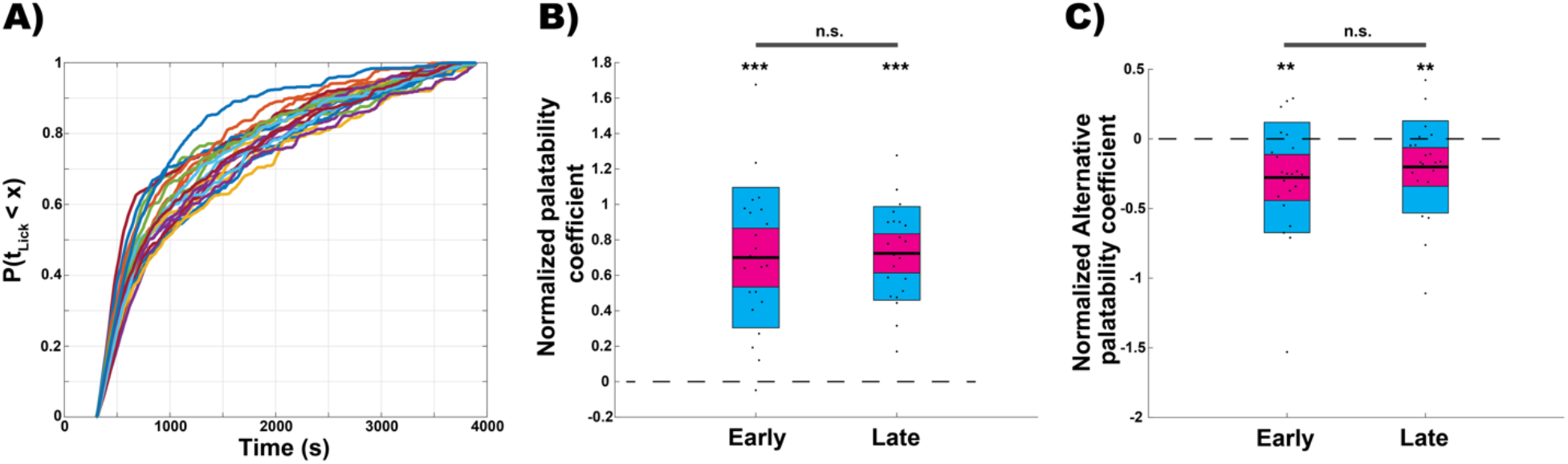
Impact of both stimuli on bout durations remains stable across a session. **A)** Cumulative distribution of lick times. Rats continue to lick throughout the session but less over time. These CDFs were used to distinguish early and late bouts for each rat. **B)** Normalized regression coefficients for current solution palatability for bouts in the early or late portion of the session. Current palatability coefficients were significantly positive for both the early (right-tailed: z = 4.06, p = 2.47e-5) and late (right-tailed: z = 4.09, p = 2.15e-5) portions of the task. Coefficients were not significantly different across portions of the session (paired: z = −.11, p = .91). **C)** Same as (B) but for the alternative solution’s palatability. Normalized coefficients were significantly negative for both early (left-tailed: z = −2.99, p = 1.4e-3) and late portions of the session (left-tailed: z = −2.76, p = 2.9e-3). Coefficients were not significantly different across portions of the session (paired: z = −.76, p = .45).

We next split bouts into those following stay decisions and those following switch decisions to ascertain whether a repeated bout at a stimulus impacted response to that stimulus or memory of the alternative stimulus. We again repeated the multilinear regression analysis on these groups individually (stay: mean ± SE *R^2^* = 0.72 ± 0.04, switch: mean ± SE *R^2^* = 0.62 ± 0.043). We found that regression coefficients for current palatability are similarly positive following a stay decision (z = 4.09, p = 2.15e-5, mean ± SE = 15.46 ±1.93) and switches (z = 4.06, p = 2.4e-5, mean ± SE = 16.04 ±2.38), with no significant difference between the two groups (z = .011, p = .91, Figure 4A). In contrast, there is a significant difference in alternative palatability coefficients in the post-stay vs. post-switch bouts: coefficients for the post-switch bouts were significantly more negative (z = 4.06, p = 2.4e-5) than those following a stay decision, which were themselves not significantly different from zero (z = −1.7, p = .088, mean ± SE = −1.09 ±1.61, Figure 4B). This result suggests that information regarding the alternative solution may only factor into decisions about sampling times following a switch between the two samples—that is, the palatability of a sampled taste, as inferred from bout durations, is impacted by the palatability of a different taste only if it (the alternative) has just been sampled.

**Figure 4.**
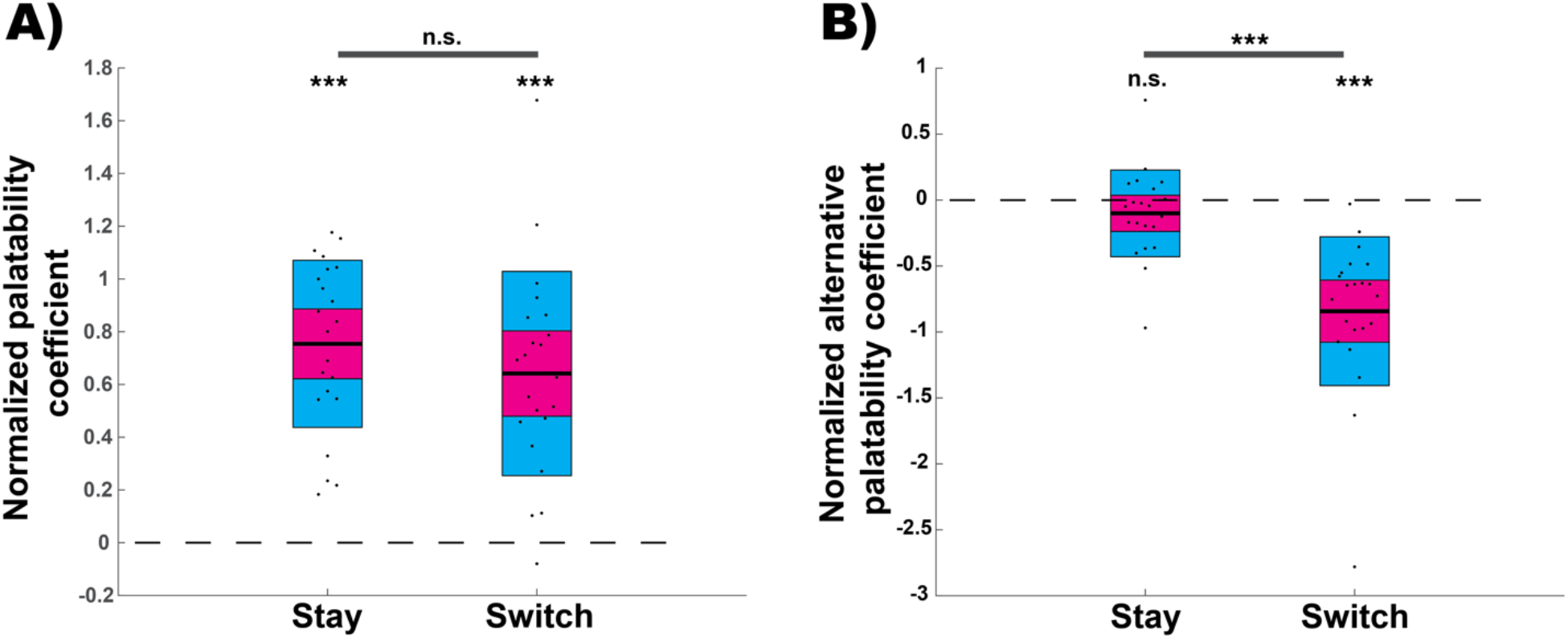
The impact on bout duration of the alternative stimulus but not the current stimulus decays following a stay decision. **A)** Normalized multilinear regression coefficients for the currently sampled solution’s palatability are significantly positive following both a stay (right-tailed: z = 4.09, p = 2.1e-5) and switch decision (right-tailed: z = 4.06, p = 2.4e-5). Coefficients were not significantly different across stay/switch conditions (paired: z = .011, p = .91). **B)** Normalized multilinear regression coefficients for the alternative solution’s palatability are not significantly different from zero following a stay decision (two-tailed: z = −1.7, p = .088) but are significantly negative for bouts following a switch decision (left-tailed: z = −4.09, p = 2.1e-5). Coefficients for bouts following a switch decision are significantly more negative than those for bouts following a stay decision (paired right-tailed: z = 4.06, p = 2.5e-5).

### Lack of dependence of results on bout definition criteria

For the analyses described above, we defined a ‘licking bout’ as sequences of spout contacts for which no interval between contacts was more than 2s (see methods). An inter-lick-interval of 2s was used as rats never switched between solutions in <2s. Of course, this is only one dividing line that could be used. Prior studies of licking microstructure in rats (Davis, 1996; Davis & Smith, 1992) have grouped licks into ‘bursts’ or ‘clusters’ based on <250ms or >500ms interlick-interval (ILI) criteria. To test that the results presented above are not artifacts of our choice of bout definition, we repeated all the above analyses using a 200ms ILI criterion (all bouts: mean ± SE R^2^ = 0.79 ± 0.033, early: mean ± SE *R^2^* = 0.72 ± 0.05, late: mean ± SE *R^2^* = 0.79 ± 0.029, stay: mean ± SE *R^2^* = 0.79 ± 0.35, switch: mean ± SE *R^2^* = 0.6 ± 0.05). In this reanalysis, the magnitudes of the resulting regression coefficients are much smaller, since bout lengths themselves are much shorter (Supplementary Figure S2). Nonetheless, all the qualitative results presented above hold (Figure 5): coefficients for current palatability are significantly positive for early vs. late (Supplementary Figure 4A) and stay vs. switch (Figure 5B) bouts, and coefficients for alternative palatability in early and late bouts do not differ (Supplementary Figure 4B); while coefficients for alternative palatability are significantly negative following a stay decision using this bout criterion (z = −3.02, p = .0012, mean ± SE = −.61 ± .19), they are again significantly more negative (mean ± SE = −2.6 ± .68) following a switch decision (z = 2.59, p = .0047, Figure 5C).

**Figure 5.**
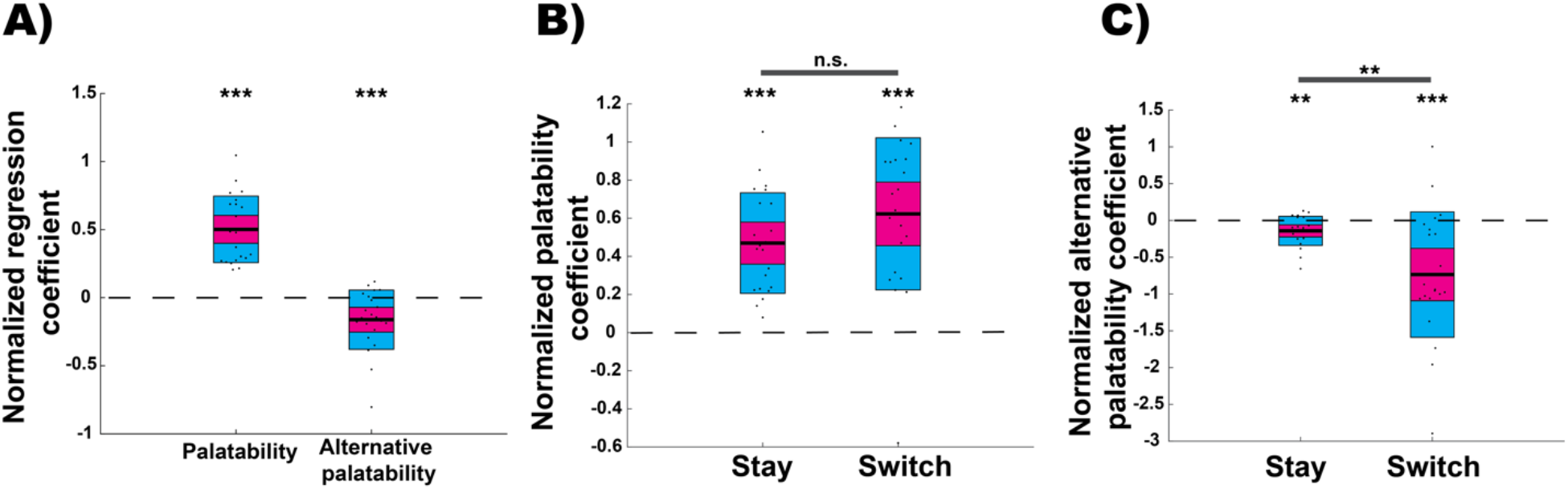
Use of a 200ms ILI interval criterion to define bouts does not qualitatively alter the results. **A)** As in Figure 2B, normalized multilinear regression coefficients for predicting bout duration using the currently sampled and alternative solutions’ palatabilities as factors are shown. Current palatability coefficients were significantly positive (right-tailed: z = 4.09, p = 2.1e-5) and alternative palatability coefficients were significantly negative (left-tailed: z = −3.15, p = 8.2e-4). **B)** As in Figure 4B, normalized multilinear regression coefficients for the currently sampled solution’s palatability are significantly and similarly positive following stay (right-tailed: z = 4.09, p = 2.1e-5) and switch decisions (right-tailed: z = 3.8, p = 7.3e-5). Coefficients were not significantly different across stay/switch conditions (paired two-tailed: z = −1.7, p = .089). **C)** Multilinear regression coefficients for the alternative solution’s palatability are shown for regressions predicting bout durations following a stay or switch decision. Using this criterion, regression coefficients for bouts following a stay decision are significantly negative (left-tailed: z = −3.08, p = .001) and following a switch decision (left-tailed: z = −3.25, p = 5.8e-4). Coefficients for bouts following a switch decision are significantly more negative than those for bouts following a stay decision (paired right-tailed: z = 2.82, p = .0024).

One difference did arise with this more stringent bout length criterion, as revealed in a comparison of Figure 5C and Figure 4B. The more stringent criterion split many prior single bouts into multiple bouts of shorter duration. The shorter duration of bouts meant that the time passed from a sampling of the alternative stimulus would often be less than previously for a repeated bout of sampling at a stimulus – that is a bout of sampling following a “Stay” decision. As a result, in Figure 5C we see a small significant impact of the alternative stimulus following a “Stay” decision that was absent in Figure 4B where bout durations were longer. Such a finding is consistent with our model in which the impact of the alternative stimulus on a current bout’s duration decays gradually over a period of many seconds after leaving that stimulus.

### Indifference of results to change in rank order of palatability

As noted above, our calculation of palatability incorporated data from days in which identical solutions were available at both spouts (see methods). Using this method, 0.1M NaCl was found to be significantly more palatable than 0.01M (Figure 1E). However, on days in which 0.1M NaCl was paired with 0.01M, rats licked more—on average, 1.3x as much—for 0.01M than for 0.1M. That is, the 0.01M solution seemed more palatable than the 0.1M solution in direct comparisons.

Regardless, we find that the results for the alternative palatability regression coefficients do not change when the palatability of the 0.01M solution is re-defined to be 1.3 times greater than that of the 0.1M solution (all bouts: mean ± SE *R^2^* = 0.8 ± 0.026, stay: mean ± SE *R^2^* = 0.61 ± 0.056, switch: mean ± SE *R^2^* = 0.76 ± 0.04): coefficients for alternative palatability remain significantly negative (z = −3.3, p = 4.5e-4, mean ± SE = −5.69 ±1.32, Figure 6A), coefficients for bouts following a switch decision remain significantly more negative than for those following a stay decision (z = 4.09, p = 2.15e-5, Figure 6C), and coefficients are not significantly different between early and late portions of the session (z = −.5, p = .61, Supplementary Figure 5); that is, our qualitative results are robust to whether the 0.01M or 0.1M solution is the more palatable and all conclusions arise from those two solutions being more palatable than the 1M NaCl solution.

**Figure 6.**
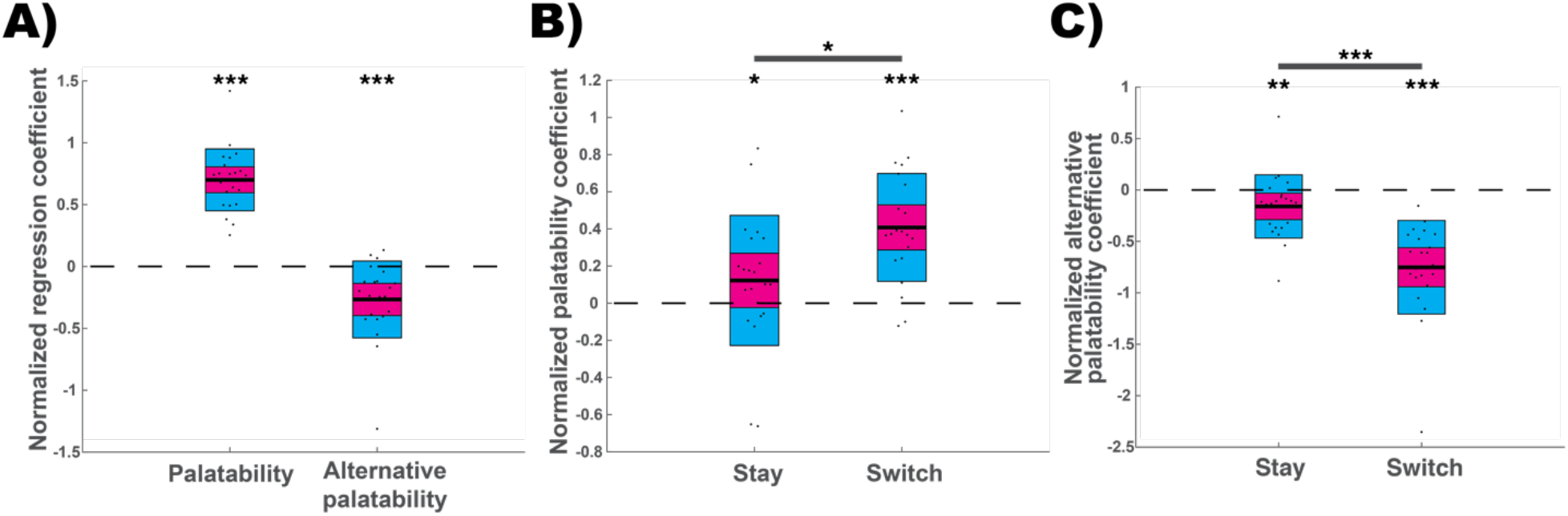
Impact of current and alternative palatability on bout duration does not depend on the relative palatability of 0.01M and 0.1M NaCl. **A-C)** The palatability of 0.01M NaCl is defined as 1.3 times the palatability of 0.1M NaCl, in accordance with relative amounts consumed in direct pairings. **A)** As in Figure 3B, multilinear regression coefficients for predicting bout duration using the currently sampled and alternative solutions’ palatabilities as factors are shown. Current palatability coefficients were significantly positive (right-tailed: z = 4.09, p = 2.1e-5) and alternative palatability coefficients were significantly negative (left-tailed: z = −3.57, p = 1.7e-4). **B)** As in Figure 5A, multilinear regression coefficients for the currently sampled solution’s palatability are significantly positive following both a stay (right-tailed: z = 2.17, p = .0148) and switch decision (right-tailed: z = 3.9, p = 4.9e-5). With these artificially altered palatabilities, the coefficients for bouts following a switch decision were significantly more positive than those for bouts following a stay decision (paired left-tailed: z = −2.2, p = .014). **C)** As in Figure 5B, multilinear regression coefficients for the alternative solution’s palatability are shown for models predicting bout durations following a stay or switch decision. With the artificially altered palatabilities, regression coefficients for bouts following a stay decision (left-tailed: z = −2.56, p = .0052) and following a switch decision (left-tailed: z = −4.09, p = 2.15e-5) are significantly negative. Coefficients for bouts following a switch decision are significantly more negative than those for bouts following a stay decision (paired right-tailed: z =4.09, p = 2.15e-5).

There are however some minor differences regarding the coefficients for current palatability. Coefficients for current palatability are significantly more positive following a switch decision (z = −3.05, p = .0011, Figure 6B) and regression coefficients of normalized bout durations in the late portion of the session were significantly smaller (less positive) than those in the early portion of the session (z = 3.18, p = 7.3e-4, Supplementary Figure 5A). These small differences, arising from a switch in ranking of the 0.01M and 0.1M solutions likely arise to changes in the importance of slaking thirst versus and balancing overall salt intake in different situations.

### Impact of palatability on transition probability

Thus far, our results describe sampling duration as a function of the palatabilities of the two available solutions, clearly demonstrating the importance of context and time on consumption choices. To understand the impact of stimulus properties on choice dynamics more fully, we analyzed the second behavioral feature determining overall consumption, namely how often a particular stimulus is visited. Deciding to visit on delivery spout instead of the other is a discrete choice that can be quantified within a transition matrix, which indicates the probability, following a pause in licking, of whether an animal returns to the same solution or switches to the alternative. We asked whether the transition probabilities between the solutions are impacted by the palatability of the current selection, that of the alternative solution, or both.

We specifically compared the transition probabilities between pairs of solutions with either a common source (e.g. 0.01M −> 0.1M and 0.01M −> 1M) or a common target (e.g. 0.01M −> 0.1M and 1M −> 0.1M). This approach allowed us to dissociate multiple contributing factors: if the palatability of the current solution influences transition probability, rats should show a higher probability of switching to a common target taste from a taste with a low palatability than from a taste with a high palatability; similarly, if alternative palatability impacts transition probability, rats should show a higher probability of switching from a common source to a solution with high palatability than to a solution with lower palatability.

We found evidence for both phenomena. That is, the palatabilities of both the current and the alternative solution were found to impact the transition probabilities in the manner expected (Figure 7A-B). These results are further supported by a logistic regression fit to predict switches based on the current and alternative palatability. In the regression model, both current palatability (p = 4.89e-31, coefficient = −.948, 95% CI = [-1.1 −.78]) and alternative palatability (p = 3.15e-21, coefficient = .63, 95% CI = [.5 .76]) are found to be significant factors.

**Figure 7.**
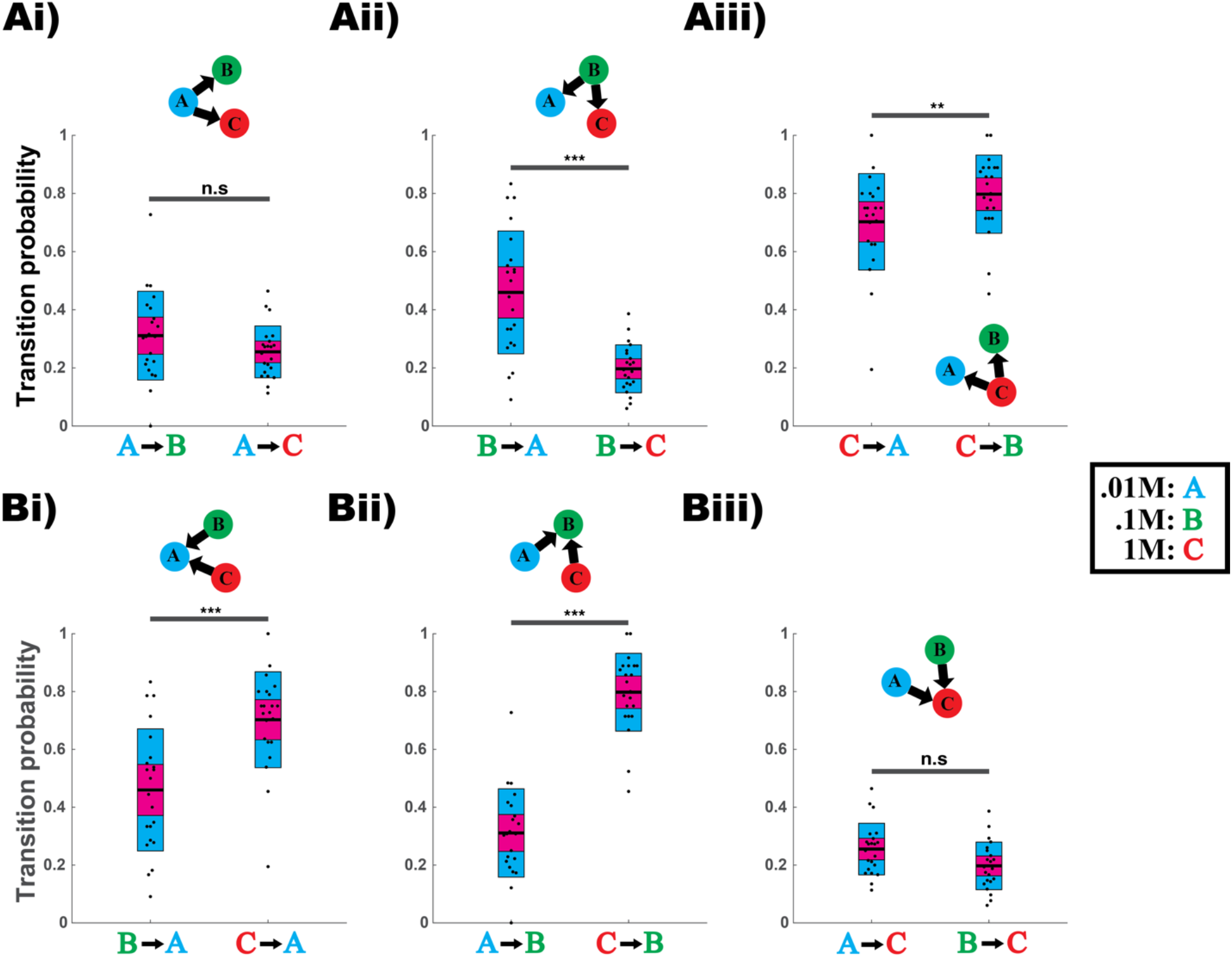
Probability of switching delivery spouts depends on the contents of the current spout and the target. **A)** Transition probabilities for transitions with a common source (0.01M −> 0.1M and 0.01M −> 1M, 0.1M −> 0.01M and 0.1M −> 1M, 1M −> 0.01M and 1M −> 0.1M) reveal the influence of a memory of the alternative solution’s palatability. **Ai)** P(0.01M −> 0.1M) and P(0.01M −> 1M) are not significantly different (paired two-tailed: z = 1.51, p = .13). **Aii)** P(0.1M −> 0.01M) is significantly higher than P(0.1M −> 1M) (paired right-tailed: z = 3.73, p = 9.4e-5). **Aiii)** P(1M −> 0.01M) is significantly lower than P(1M −> 0.1M) (paired left-tailed: z = −2.52, p = 5.8e-3). **B)** Transition probabilities for transitions with a common target (0.1M −> 0.01M and 1M −> 0.01M, 0.01M −> 0.1M and 1M −> 0.1M, 0.01M −> 1M and 0.1M −> 1M) reveal the influence of the last sampled solution’s palatability on switch probability. **Bi)** P(0.1M −> 0.01M) is significantly lower than P(1M −> 0.01M) (paired left-tailed: z = −3.63, p = 1.3e-4). **Bii)** P(0.01M −> 0.1M) is significantly lower than P(1M −> 0.1M) (paired left-tailed: z = −4.09, p = 2.15e-5). **Biii)** P(0.01M −> 1M) is significantly higher than P(0.1M −> 1M) (paired right-tailed: z = 2.43, p = 7.5e-3).

### No evidence for memory across days

The above analyses suggest, among other things, a short-lasting memory of which taste had recently been sampled. In our last set of analyses, we investigated whether rats maintained a memory of a session from a prior day and whether this biased their initial sampling choices. To do this, we counted the number of times rats first visited the side they preferred (had the most licks at) on the prior day and compared this to the number expected. Given a null hypothesis of no memory across days, the expected number is given by the binomial distribution with p = q = 0.5. Our results are consistent with the null hypothesis that rats did not carry a preference for side across days (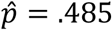, 95% *CI* = [.424 .547], Figure 8).

**Figure 8.**
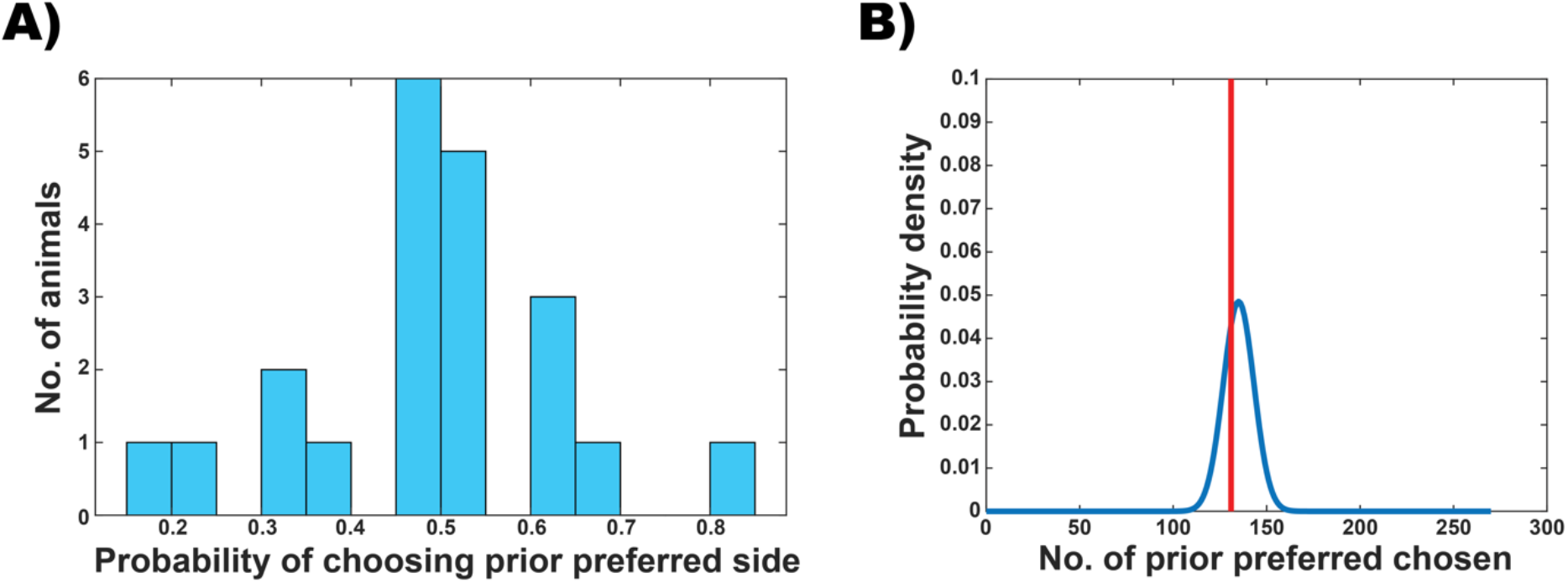
No evidence for preference across days. **A)** Histogram showing the fraction of times rats first sampled from the side they preferred on the prior day. **B)** (blue) Probability density function for the total number of times (across all rats) that rats first sampled from the side they preferred on the prior day which is given by the binomial distribution with N = 270, p = q = 0.5. (red line) Total number of times rats first sampled from the preferred side from the prior day.

### Comparison to spiking network models

In a recent modeling study (Ksander et al., 2021), we instantiated a two-pool spiking network that implements continuous-time decision-making. One pool represents the ongoing decision to continue sampling, which received greater excitatory input with increased palatability of the stimulus. The other pool represents the decision to stop sampling and consider switching to an alternative stimulus. These two pools mutually inhibit one another such that only one can be active at a time and transitions between these two states are induced by noise fluctuations. We identified the durations of periods of continuous activity of the neurons representing a continuation of sampling as the durations of sampling bouts.

We found competition in the lengths of such bout durations in response to alternating stimuli— that is, following a strong (palatable) stimulus with a long bout, the palatability response to the subsequent stimulus was relatively diminished. The competition arose from a slow synaptic depression in the model, which acted to weaken connection strengths following sufficiently long periods of high neural activity, so we hypothesized that the competition between successive stimuli would diminish over the timescale of recovery from that synaptic depression. We predicted that the impact of the alternative stimulus on bout duration would, therefore, be significantly lower, during a second or later successive bout at the same stimulus, as compared to the first bout at that stimulus following a switch, just as seen in the behavioral data (Figures. 5–7). To test this prediction, we adapted the stimulus protocol used in our prior study (which enforced strict alternation) such that following any state transition indicating the end of a bout of sampling, the subsequent stimulus presented was chosen randomly, allowing for repeated “stays” as well as “switches” between stimuli.

Our results are shown in Figure 9, for which we produced regression coefficients in the same manner as Figs. 5–7 using distinct pairs of three stimuli of different strengths, each representing different palatability. The results are qualitatively identical to the behavioral data (Figures 3 and 5–7) with the alternative stimulus having a competitive impact on bout duration (a negative regression coefficient, Figure 9A) but with the impact diminished following a repeat bout (a “stay” transition, Figure 9C) at the same stimulus.

**Figure 9.**
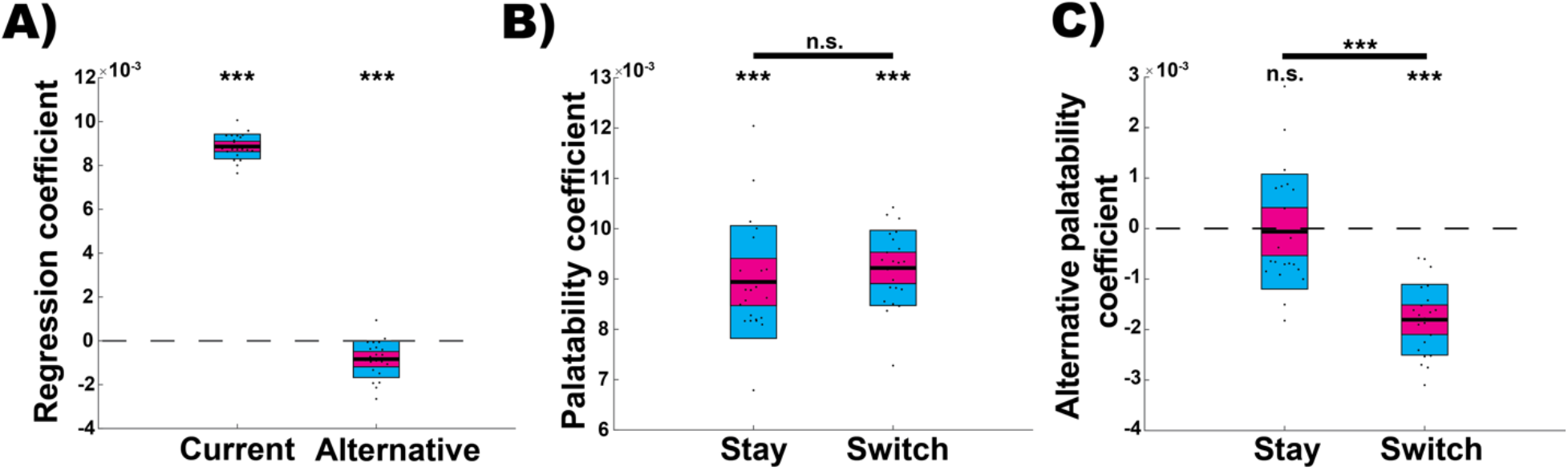
Model network replicates key features of rat behavior. **A)** As in Figure 3B regression coefficients for predicting bout duration as a function of current and alternative palatability are shown for model networks. Similar to rats, coefficients for current palatability are significantly positive (right-tailed: z = 4.09, p 2.15e-5) and coefficients for alternative palatability are significantly negative (left-tailed: z = −3.83, p = 6.38e-5). **B**) As for rats, palatability coefficients for both stay (right-tailed: z = 4.09, p = 2.15e-5) and switch (right-tailed: z = 4.09, p = 2.15e-5) bouts were significantly positive and are not significantly different between groups (paired two-tailed: z = −1.8, p = 0.07). **C**) As for rats, alternative palatability coefficients for bouts following a stay decision were not significantly different from zero (two-tailed: z = −1.28, p = 0.19) whereas coefficients for bouts following a switch decision are significantly negative (left-tailed: z = −4.06, p = 2.47e-5)).

### Diminishing impact of the alternative stimulus over time

Both the rats and spiking model display a reduced or non-existent impact of the alternative stimulus’ palatability on bout durations following a “stay” decision. In the spiking model, we can attribute this effect to the decaying impact of synaptic depression. However, for the rats, this could be due to either an effect of time (i.e. a decaying ‘memory’ trace) or the presence of a distractor. To determine which source most likely causes the rat behavioral phenomenon, we performed two additional analyses. In the first, we determined, for each bout, the amount time that had elapsed since the rat’s last visit to the alternative stimulus (Δ*t_last visit_*). We then split the bouts based on a threshold which we varied from 10s to 60s, the timescale over which synaptic depression effects might last. For each split, we performed the same multilinear regression analysis described above.

The results of this re-analysis are displayed in Figure 10, which shows that for all Δ*t_last visit_* thresholds from 10s-60s, alternative palatability regression coefficients are significantly more negative for the bouts with a smaller Δ*t_last visit_*. That is, the impact of the alternative stimulus does indeed decay over time but does not rule out a stimulus-related effect—following a “stay” decision, the immediately preceding stimulus at the same lick spout could be thought to act as a “distractor” for the memory of the stimulus at the alternative lick spout. For example, if the sequence of solutions sampled is A – B – B, the first bout at B represents a distracting stimulus when assessing the impact of A’s palatability on the second bout at B.

**Figure 10.**
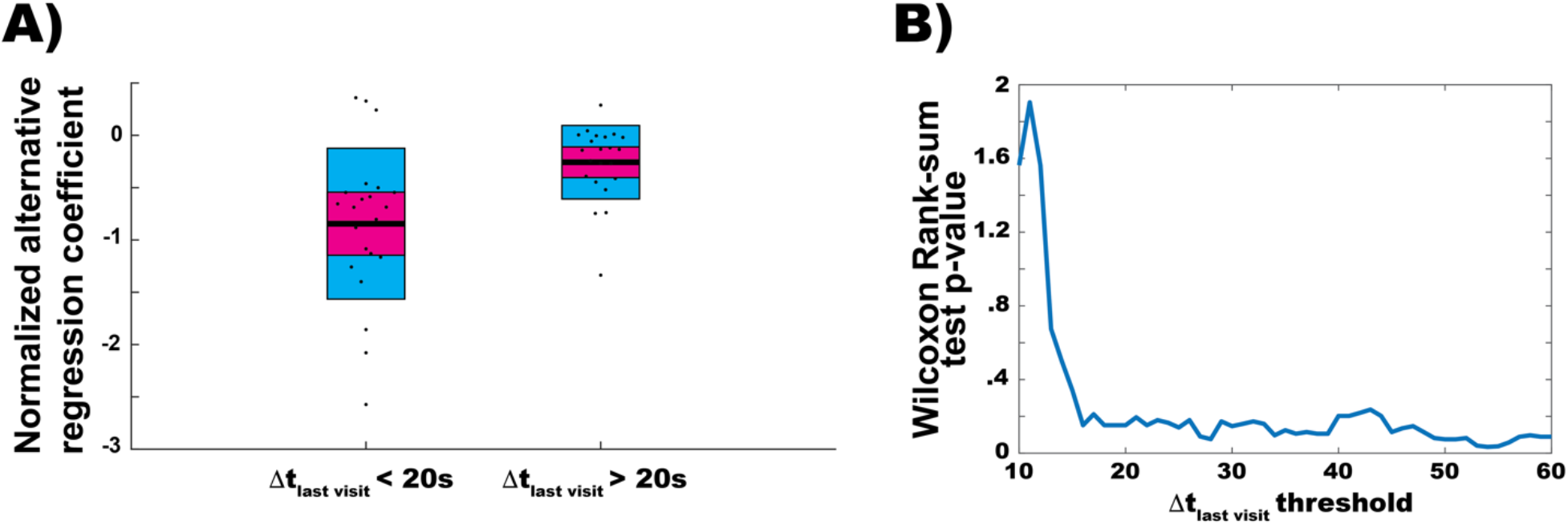
The impact of the alternative solution’s palatability decays over time. **A)** Normalized regression coefficients for the impact of the alternative solution’s palatability for bouts where Δ*t_last visit_* is less than 20s or greater than 20s. **B)** The analysis in (A) was repeated using different Δ*t_last visit_* separation thresholds (in (A) it was 20s) from 10s to 60s. The regression coefficient distributions for each threshold were compared using the Wilcoxon Rank-sum test and the resulting p-value is plotted on the y-axis.

To determine whether it is time, that distracting stimulus, or a combination of both that leads to the diminishing effect of the alternative stimulus, we performed an N-way ANOVA with the current and alternative solutions’ palatabilities, Δ*t_last visit_*, and the presence of a distractor as factors. We additionally included interaction terms for the alternative solution’s palatability and Δ*t_last visit_* as well as the alternative solution’s palatability and the presence of a distractor. Lastly, to limit the impact of the likely non-linearity of the interaction of alternative palatability and time, we restricted the ANOVA to bouts with a Δ*t_last visit_* < 60s and matched the Δ*t_last visit_* distributions for bouts with and without a distractor.

This ANOVA revealed a statistically significant interaction between alternative palatability and Δ*t_last visit_* (F(1,347) = 16.13, 1e-4) but not between alternative palatability and the presence of a distractor (F(1,347) = 1.91, p = 0.1678). Simple main effects analysis showed that current palatability, alternative palatability, and *Δt_last visit_* had a statistically significant effect on bout duration each with a p-value < 1e-4. Simple main effects analysis showed that the presence of a distractor stimulus did not have a statistically significant effect on bout duration (p = 0.0968). Based on this analysis, we conclude that the time since the last visit to the alternative stimulus is the dominant factor causing a reduced impact of alternative palatability in bouts following a stay decision. This result, interpreted through the lens of our spiking model, suggests the functioning of decay of synaptic depression processes.

## Discussion

Palatability is typically evaluated through measurement of the amount of a food or solution consumed. Given that the mean amount consumed per lick has been shown to be very stable across time (Wilcove & Allison, 1972), in this study we consider the total number of licks of a solution as a reasonable quantification of palatability. But since rats sample a solution in clearly demarcated bouts with high frequency (approximately 6 Hz) regular sampling within a bout, we have gone beyond this simple measurement, making use of two dynamic factors that have been shown to provide “real-time” measures of palatability: the duration of the bouts and the total number of bouts. In theory, bout durations could be independent of a stimulus such that its palatability is only evident in the total number of bouts, but prior work (Davis, 1996) has shown that the more palatable a stimulus, the longer the bouts, a result which we replicate here. What our work adds is an assessment of whether, in a preference test, the palatability of the alternative stimulus impacts the behavior of an animal at its current stimulus. Our main finding is that this impact is strong: the more palatable the alternative stimulus, the shorter the bouts of licking at (i.e., the less palatable is) the current stimulus.

A second factor that goes into determining the total number of licks at a spout is the number of times the spout is visited. We found that this factor as well is impacted by available alternatives—that the probability of a return increases with the palatability of the stimulus just tasted (to which it would return). Such behavior necessarily translates into more bouts at sources of high palatability, leading to fewer at an alternative. We found that even following a pause in sampling at one lick spout, memory of the alternative impacted the likelihood to repeat at the same spout. That is, the more palatable the alternative, the greater the probability the animal would transition to that alternative spout and the less likely to repeat a sample at the same spout. Thus, in contrast to bout durations, the choice of which spout to lick from is impacted by memory of the contents of both lick spouts, a memory that persists beyond the duration of and survives the interference of a single bout of sampling.

Our findings support a recent model (Ksander et al., 2021) in which the duration of a bout is given by the duration of a particular state of activity in a neural circuit. In the model, noise fluctuations terminate states of activity, leading to an exponential-like distribution of state durations, just as we find in the behavioral data (Supplementary Figures 1–3). Moreover, the impact of the neural-circuit level process of synaptic depression in the model leads to a competitive impact between successive stimuli, such that following a highly palatable stimulus a subsequent bout duration is shorter than otherwise expected. The biological processes that naturally instantiate synaptic depression have a limited timescale, such that in the model the impact of competition on bout durations diminishes over time and is much weaker for bouts following a return to a stimulus when the time passed since the visit to the alternative has increased (Figures 9–10). We find a similar effect in our behavioral data, with no impact of the alternative stimulus on durations of bouts of sampling that do not directly follow a switch from that alternative stimulus (Figure 4B). This fact provides additional validation for model and theory.

When a taste stimulus is considered palatable or unpalatable, the implicit suggestion is that palatability is a property of a substance to be ingested. However, in practice, palatability is a measure of behavior—typically the total amount of a substance consumed—so is inherently dependent on the state of an animal and the context in which the animal is sampling the stimulus. In our study, even the rank order of palatability can be altered by context: when a rat has two lick spouts available to it, both of which contain the same solution, in accordance with prior work (Sadacca et al., 2012) rats lick more for 0.1M NaCl than 0.01M NaCl (Figure 1), suggesting that 0.1M NaCl is more palatable to rats than is 0.01M NaCl; when one lick spout contains 0.1M NaCl and the other contains 0.01M NaCl, however, rats lick more often at the spout containing 0.01M NaCl, suggesting a context-dependent switch in relative palatability and preference of the two salt solutions. Such a switch could arise from a trade-off between the competing needs of slaking thirst and balancing sodium levels, with most preferrable concentrations depending on what else is available and amount already drank in a session. Regardless, however, our findings in this paper regarding the interactions between stimuli were robust to this switch. That is, we could quantify the palatability of 0.1M NaCl and 0.01M NaCl using the results of sessions when they were paired with each other rather than on the sessions with a single stimulus, without altering any of our findings on how the palatability of the alternative stimulus impacts the behavior at a lick spout.

Our findings of a competitive interaction between the palatability—or value—of simultaneously available stimuli, combined with our model of the process (Ksander et al., 2021), add richness to the foraging literature, in which behavior is discussed historically in terms of the Marginal Value Theorem (Charnov, 1976). The theorem prescribes optimal behavior in an environment with multiple sources, at each of which the rate of reward diminishes with the time an animal spends at the source. Specifically, an animal should only stay at a food source until its rate of reward has depleted to the mean rate of reward it would achieve by moving from alternative source to alternative source and remaining the optimal time at the alternative sources. Our behavioral findings and model are in qualitative accordance with such behavior in that the more palatable an alternative (*i.e*., the greater the mean rate of reward in the environment) the less time spent at a source. Moreover, such reduction is ameliorated over time, an effect also in accordance with optimal theory, as the greater the time spent between sources while foraging, the lower the mean rate of reward, so time spent at a diminishing source increases.

Of course, it is unlikely that our relatively simple network model encapsulates the complexities underlying time valuation in naturalistic foraging. Importantly, unlike in foraging studies, in preference tests the potential rate of reward at a lick spout is constant, so if one spout contains more rewarding solution than the other, the optimal behavior of an animal would be to stay at the more rewarding spout as soon as it has sampled both. That the animals do not behave in such a manner, but continue to sample even aversive stimuli many times, is either an indication of limited memory duration (*i.e.,* they forget what is in each spout) or a strong drive to explore in case the environment changes.

## Methods

### Preregistration

This study was not preregistered.

### Subjects

22 adult, taste-naïve, Long-Evans rats (14 female, 8 male) from Charles River were subjects. All subjects were individually housed and kept on a 12/12 hr light/dark cycle. Experiments were conducted during the light cycle and complied with Brandeis University Institutional Animal Care and Use Committee guidelines.

#### Behavioral apparatus

The preference test was carried out in 1’ x 1’ x 1’ custom acrylic chambers. Each chamber has 3 holes through which rats could lick a stainless-steel delivery spout. There is one hole on each of the left, right, and back walls of the chamber. For this study, only the left and right sides ever had a delivery spout. In order to record licks, a custom circuit, based on a published design (Hayar et al., 2006) was used. A small voltage was applied to the stainless-steel floor of the chamber such that when the rat licked one of the spouts, a voltage deflection (measuring the water-metal junction potential) was recorded. A RaspberryPi was used to both supply power to the floor and record licks using custom Python software.

### Preference test

Rats were water deprived for 22 hours prior to the first habituation session. The preference test timeline consisted of 16 one-hour sessions of which the first 2 were habituation sessions with only one bottle of distilled water (dH_2_O) available on one side of the experimental chamber (the side was switched for the second habituation session). Following each session, rats were given one hour of ad lib access to water in their home cage such that they were deprived of water for 22 hours prior to each session. After the two habituation days, the first day of the preference test was always a session with two bottles of dH_2_O. This was followed by six consecutive days of pairings of three NaCl concentrations (0.01M, 0.1M, 1M), with all combinations including self-pairings used, one pairing per day. This was then repeated for a second week such that each rat experienced two dH_2_O only sessions and two pairings of each combination of NaCl concentrations. These concentrations were used because they had been previously measured to have different palatabilities (Sadacca et al., 2012) covering both palatable and unpalatable (relative to water) tastants.

### Lick identification

Licks were identified via a semi-automated process using custom MATLAB software. A simple threshold could not be used to identify licks because both licks and nose pokes were picked up as large voltage deflections. Additionally, occasionally a rat would maintain contact with the lick spout while licking, resulting in a sustained voltage deflection on top of which licks could be identified. As a result, we produced a dataset of hand-identified licks from the data of the first few subjects and used MATLAB’s neural network toolbox to train a bidirectional LSTM recurrent neural network to predict the presence or absence of a lick at any point in time. These automatic identifiers were then used as a first pass on all future data to capture presumptive licks, which were then accepted/discarded by eye based on the stereotypical shape and timing of licks. Lastly, a final pass over the data was made by eye to ensure that no licks were missed by the neural network.

#### Lick bout identification

Following identification, licks were grouped together into ‘bouts’ based on 2 different inter-lick interval (ILI) criteria. That is, we repeated all analyses described using bouts defined by 2 different ILI criteria to determine how our results depended on this somewhat subjective threshold. Based on previous studies of licking dynamics in rats (Davis, 1996) and our own investigation of ILI distributions, we grouped together licks with 200ms ILIs into lick ‘bouts’ (also referred to as lick clusters). We also used a more nuanced criterion which we believe better represents active engagement with a lick spout (indicating an ongoing ‘stay’ decision). This criterion consisted of grouping together adjacent licks in which there was no period >2s between them in which there was no activity on the recording channel. This means that if the rat nose-poked the solution spout in between licks such that the ILI was >2s but there was intervening activity on the channel such that there was no period of >2s of silence, then these licks would be grouped together. We included this criterion since we are primarily concerned in this study with the rats’ decisions to leave a solution spout and not on the microstructure of their licking behavior.

### Measurement of palatability

To measure the palatability of each concentration of NaCl, we analyzed data exclusively from sessions where a solution was paired with itself. This was done to separate the data used to compute palatabilities from those used in the multilinear regressions. While the measurements of palatability are derived from lick counts and the multilinear regressions predict bout duration, these two variables are correlated and we therefore wanted to prevent a situation in which the outputs of the regressions were in part derived from the same data as the inputs to the regression model. The palatabilities were defined relative to water such that the relative palatability of solution X was:

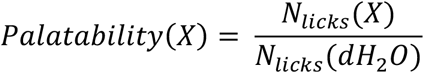

where *N_licks_(X)* is the total number of licks to tastant X across both sessions when X was paired with itself. *N_licks_(dH_2_O)* is the same except for dH_2_O only sessions.

### Linear and logistic regression models

To assess the differential impacts of the palatability of the currently sampled solution and alternative solution on the current bout duration, we performed multilinear regressions using MATLAB’s *regress* function to predict bout duration with palatability of each solution as factors. This was done for all bouts together, as well as for subsets of bouts depending on if they were ‘early’ or ‘late’ in a session, or following a stay or switch decision. Whenever ‘normalized regression coefficients’ are presented, the normalization results from dividing the resulting regression coefficients by the mean bout duration of the group of bouts from which the coefficients were determined.

As part of the ANOVA analysis determining the effect of distractor stimuli, the distributions of times since leaving the alternative spout (Δ*t_last visit_*) were matched for the groups of bouts with and without a distractor stimuli. This matching was performed using the ***matchpairs*** MATLAB function with the pairwise difference matrix of Δ*t_last visit_* between the two groups as input and a ‘costUnmatched’ parameter of .1.

As one method of measuring the impact of current/alternative palatability on switch probability, we performed logistic regression using MATLAB’s ***fitglm*** function to predict a switch (0 or 1) with the last sampled solution’s and alternative solution’s palatabilities as factors.

### Separating early and late bouts

We separated bouts for each session on a per-animal basis into ‘early’ or ‘late’ bouts by analyzing the 2^nd^ derivative of the cumulative distribution of lick times across all sessions. First, a smoothed probability density function of lick onset times was computed using MATLAB’s ***ksdensity*** function with a bandwidth of 200s (controlling the amount of smoothing). The cumulative density function of this pdf was then computed and its 2^nd^ derivatives approximated. The time point with the minimum 2^nd^ derivative was then used as the divider between early and late bouts.

### Statistical tests

Unless otherwise stated, all z and p-values reported in this paper are from the Wilcoxon signed rank test performed using MATLAB’s ***signrank*** function. Tests of whether the median of a distribution is significantly positive/negative utilized the right/left-tailed test respectively. Tests of differences between distributions were done using a paired test where data points were paired by animal or, in the case of the spiking model, points were paired by network. Additionally, all tests had an N = 22.

#### Simulation protocol

Simulations were carried out using a recently published model (Ksander et al., 2021). In brief, leaky integrate-and-fire neurons were designated excitatory or inhibitory and assigned either to a group whose activity promoted a continuous decision of “Sample” at the current stimulus (termed “Stay” in the prior paper) or a group whose activity promoted a decision to “Leave” the stimulus (termed “Switch” in the prior paper). In the original paper we had assumed that leaving one stimulus meant the animal always switched to the alternative. For this paper, we produced new simulations with successive stimuli chosen randomly (from the two in use for a particular session) with equal probability, to investigate consequences of a “Return” to the same stimulus following a “Leave” decision that ended a bout.

Connections between types of neurons were arranged in a manner of self-excitation and cross-inhibition such that activity of one type of neurons *(e.g.,* representing continuous sampling) could maintain itself while at the same time suppressing activity of the other type of neurons *(e.g.,* representing “Leave”). Activity of the “Sample” neurons while suppressing the “Leave” neurons would represent a state in which the animal continues to sample a stimulus. Noise fluctuations would irregularly cause a transition from such a “Sample” state to a “Leave” state. We would ensure the “Leave” state was transient by reactivating “Sample” neurons to represent the animal’s commencement of the next sampling bout. As in our behavioral data, such noise-induced transitions to terminate a bout of sampling resulted in an exponential-like distribution of bout durations.

We assessed two types of model, in one type, the “entice-to-stay” model, the mean bout durations were determined by stimulus-dependent input to excitatory neurons in the “Stay” pool, such that the more palatable the represented stimulus, the greater the input. In the other type, the “repel-to-leave” model, the mean bout durations were determined by stimulus-dependent input to excitatory neurons in the “Leave” pool, such that the more palatable the represented stimulus, the lower the input. We also test both types of model in this work.

All synapses in the model include synaptic depression, comprising a fast (300 ms) process representing docking of new vesicles after vesicle release and a slow (7 sec) process representing replenishment of a reserve pool of vesicles. Synaptic depression is key in producing the competitive interaction across time as after a period of strong activity the connections that sustain activity are weakened, impacting the response of the network to a subsequent stimulus, until recovery of the supply of vesicles is complete.

### Properties of model neurons

Individual neurons were simulated with an exponential leaky integrate-and-fire model (Fourcaud-Trocme et al., 2003) following the equation:

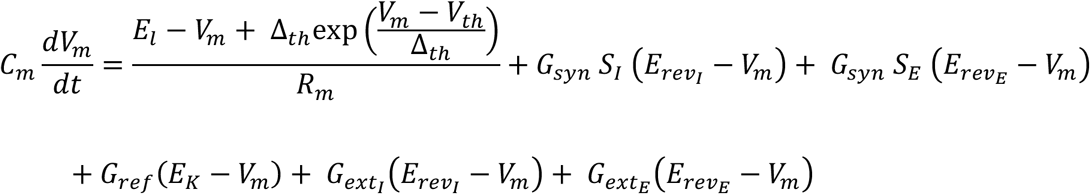

where *V_m_* is the membrane potential, *C_m_* is the total membrane capacitance, *E_l_* is the leak potential, *R_m_* is the total membrane resistance, *Δ_th_* is the spiking range, *V_th_* is the spiking threshold, *S* is the synaptic input variable, *G_syn_* and *E_rev_* are the maximal conductance and reversal potential for synaptic connections, *G_ref_* is the dynamic refractory conductance, *E_K_* is the potassium reversal potential, and *G_ext_* is the input conductance. The “E” and “I” subscripts denote the variables specific to excitatory and inhibitory channels, respectively (e.g. *S_E_* and *E_rev_E__* are the synaptic input and reversal variables for excitatory channels; *S_I_* and *E_rev_I__* are the corresponding inhibitory variables). This equation simulates the neuron’s membrane potential until *V_m_ > V_spike_*, at which point the neuron spikes.

When a neuron spikes, *V_m_* is set to the *V_reset_* value. Additionally, the neuron’s refractory conductance, synaptic output, *s*, and synaptic depression (noted as *D*) are updated according to the equations:

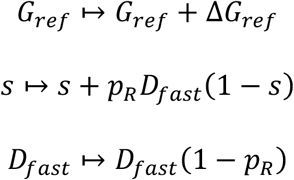

where *ΔG_ref_* is the increase in refractory conductance, and *p_R_* is the vesicle release probability following a spike.

In the timestep immediately following a spike, the neuron’s membrane potential continues to follow the exponential leaky integrate-and-fire model equation. In this equation the separate excitatory (*S_E,i_*) and inhibitory (*S_I,i_*) synaptic inputs for cell *i* are obtained from the sum of all presynaptic outputs multiplied by the corresponding connection strengths, *W_ij_*, from neurons *j* (see *Network architecture and connections)*:

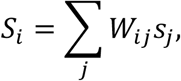

each of which decay with the appropriate (excitatory or inhibitory) synaptic gating time constant *τ_s_* according to:

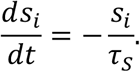

Likewise, refractory conductance decays with the time constant *τ_ref_* according to:

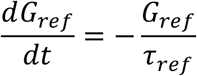

The *G_ext_* input conductance serves as both noisy-background and stimulus inputs in the same manner. Inputs were modeled as Poisson spike trains with rates *r_noise_* and *r_stimulus_*, which produce input spikes (from all sources) at timepoints *{t_sp_}*. Please note, the noisy-background includes both excitatory and inhibitory spiking input (included in *G_ext_I__* and *G_ext_E__*, respectively); the *r_noise_* parameter specifies the rate for both excitatory and inhibitory background noise. The input conductance values for a given timepoint, *t*, are updated as:

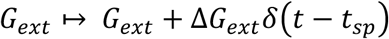

where the conductance increases by *ΔG_ext_* at the time of each input spike. The input conductance otherwise decays with the time constant *τ_ext_* according to:

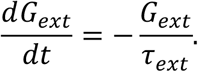

The cellular parameters with values specific to excitatory neurons (e.g. that differ from inhibitory values) are: *E_rev_E__* = 0 *mV*, *τ_s_* = 50 *ms*, and *τ_ext_* = 3.5 *ms*. The complementary values for inhibitory neurons are: *E_rev_I__* = −70 mV, *τ_s_* = 10 *ms*, and *τ_ext_* = 2 *ms*. The remaining parameters applicable to both excitatory and inhibitory neurons are: *G_syn_* = 10 *nS*, *p_R_* = .1, *τ_fast_* = ^300^*ms*, *t_slow_* = 7*S*, *P_slow_* = .5, *E_l_* = 70*mV*, *E_K_* = 80*mV*, *V_reset_* = 80*mV*, *R_m_* = 100 *M⋂*, *C_m_* = 100 *pF*, *V_spike_* = 20*mV*, Δ*G_ext_* = 1 *nS*, *V_th_* = −50 *mV*, Δ*_th_* = 2 *mV*, *τ_ref_* = 25 *ms*, and Δ*G_ref_* = 12.5 *nS* . The Poisson spike-train parameters *r_noise_* and *r_stimulus_* are described in the next section. Neurons were simulated with a simulation timestep *dt* = .1 *ms*.

### Synaptic depression

We modeled synaptic depression using two separate timescales, noted in the previous spike-update equations as *D_slow_* and *D_fast_*. These two variables reflect, respectively, the fraction of the maximum number of vesicles available in the reserve pool and the release-ready pool. Following a spike, the variables recover to a value of one with different timescales, because vesicles regenerate and are replenished slowly in the reserve pool, but may dock and become release-ready much more quickly once available in the reserve pool

Specifically, *D_slow_* represents the ratio of currently available reserve-pool vesicles out of the maximum possible, that is 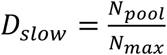. These dock quickly at empty docking sites on the timescale *τ_fast_*, but are replaced slowly on the timescale *τ_slow_*. *D_fast_* represents the ratio of docked vesicles out of total docking sites, that is 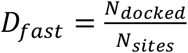. We also incorporate the constant parameter, *f_D_* = 0.05, which is equal to the ratio of the number of docking sites to the maximum size of the reserve pool of vesicles, 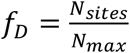. Only docked vesicles can be released immediately following a spike, such that upon each spike we update *D_fast_* ↦ *D_fast_* (1 – *p_R_*) where *p_R_* is the vesicle release probability.

During sustained spiking, the fast-docking can maintain a firing-rate dependent supply of docked vesicles until the reserve pool depletes. Vesicles dock at empty sites according to:

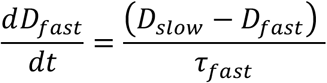

Reserve-pool vesicles fill the empty docking sites on the fast timescale *τ_fast_*. On the other hand, the reserve-pool regenerates much more slowly according to:

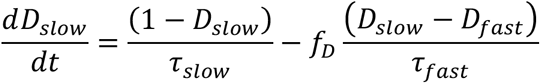

The first term represents the reserve-pool vesicle regeneration on timescale *τ_slow_*. The second term 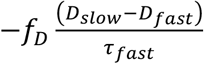 accounts for the vesicles lost due to docking.

Our model reflects the empirical evidence showing the effects of synaptic-depression at short timescales on the order of milliseconds, and longer timescales on the order of seconds (Abbott et al., 1997; Varela et al., 1997); depression timescales on the order of minutes have even reported in non-mammalian animals (Tabak et al., 2000). Additional, recent evidence (Kusick et al., 2020) directly supports our fast-depression mechanism where available vesicles quickly refill empty docking sites. Our model provides a coherent mechanism for both fast-acting and long-lasting synaptic depression effects.

#### Network architecture and connections

Each network consists of 250 individual neurons, split into two populations of 100 excitatory cells (*i. e.,* “stay” and “switch” populations, E_stay_ and E_switch_) and two populations of 25 inhibitory cells (I_stay_ and I_switch_). For each pair of connected populations (or for self-connected excitatory populations) pairs of cells were connected probabilistically with a probability, *P*(*connection*) = .5. The strength of connections was symmetric across “stay” and “switch” populations but depended on whether presynaptic or postsynaptic cells were excitatory or inhibitory. The connection strengths used for this paper are given by the open (‘repel-to-leave’) and closed (‘entice-to-stay’) squares in Figure 2 of (Ksander et al., 2021). That is, for entice-to-stay networks, E-to-I connections had a fixed weight of 0.0909 and I-to-E connections had a fixed weight of 9.6192. Repel-to-leave networks and a fixed E-to-I weight of 0.4242 and a fixed I-to-E weight of 9.4939.

### Network states and stimuli

A network’s active state was evaluated by comparing the mean values of synaptic output, *s_E_*, averaged across all excitatory cells in each of the two excitatory populations. Specifically, when the difference between the mean output of the previously less active excitatory population exceeded that of the previously more active excitatory population by a threshold of 0.02 consistently for 50ms, we recorded a state change.

We did not simulate the animal’s behavior in between bouts of sampling a stimulus. Once the excitatory neurons in the “switch” population (E-switch cells) were recorded as more active than those in the “stay” population, using the threshold mentioned above, we removed the stimulus input to the network. 100 ms later, we induced a subsequent transition back to the “stay” state to represent the animal initiating a new bout of stimulus sampling. The transition back to sampling was accomplished by halving the noisy background input to E-switch cells until the network transitioned again to the “stay” state. At all other times in simulations, the noisy background input remained constant. Once a transition to the “stay” state was recorded (by excitatory cells in the “stay” population being more active than those in the “switch” population) input stimulus was applied to indicate the next bout of sampling. The choice of subsequent next stimulus was probabilistic, with 50% probability of each of the pair being compared in the simulated preference test. Individual taste preference task simulations lasted 1500 seconds total. Each simulation compared sampling bout durations in response to two stimuli each with a fixed value across the session. For a given network the stimulus inputs targeted the same population for all sessions.

To produce linear regression coefficients in Figure 10, we regressed the log of the state duration as a function of the stimulus strengths used, because state durations depend exponentially on stimulus strengths in our model, which is based on noise-induced transitions between attractor states (Kramers, 1940; Miller & Wang, 2006).

## Data Availability

Data and analytical codes can be found via https://github.com/benballintyn/behavior. Simulation codes can be found via https://github.com/johnksander/naturalistic_decision_making.

## Supplementary figures

**Supplementary Figure 1.**
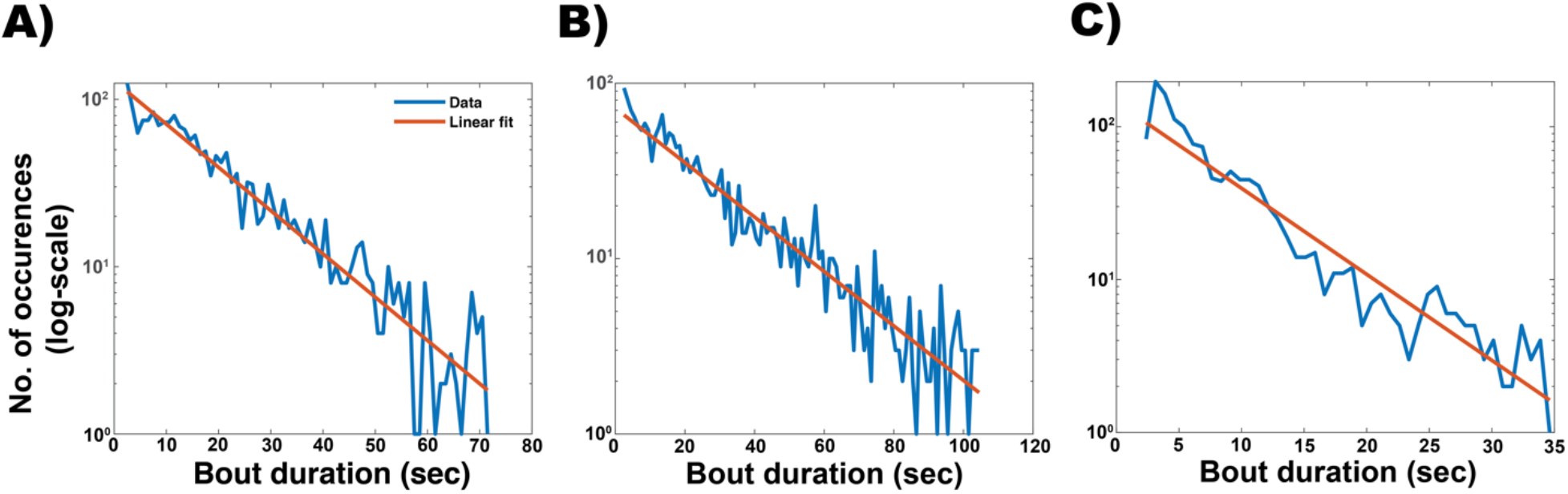
Bout durations are roughly exponential. **A)** Bout duration frequencies for the 0.1M NaCl are plotted on a log-scale with an exponential fit shown (orange). **B)** Same as (A) but for bouts at the 0.1M NaCl. **C)** Same as (A,B) but for bouts at the 1M NaCl.

**Supplementary Figure 2.**
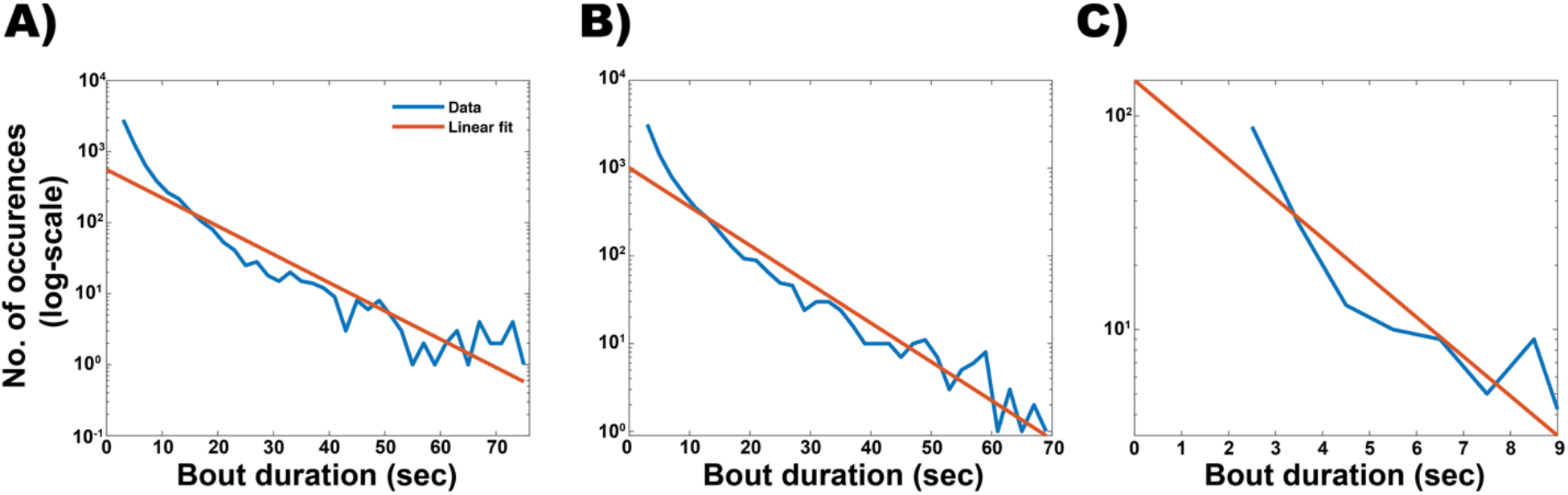
Bout durations using the 200ms ILI bout definition are roughly exponential. **A)** Bout duration frequencies for the 0.1M NaCl are plotted on a log-scale with an exponential fit shown (orange). **B)** Same as (A) but for bouts at the 0.1M NaCl. **C)** Same as (A,B) but for bouts at the 1M NaCl.

**Supplementary Figure 3.**
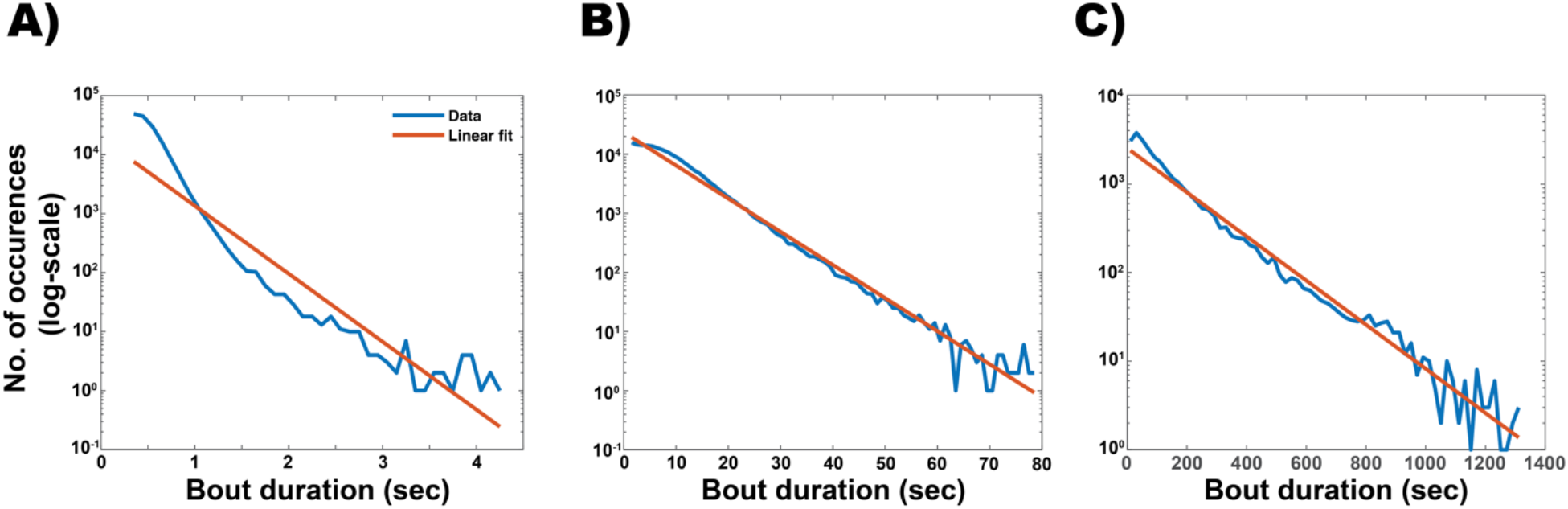
Bout durations produced by the spiking model are roughly exponential. **A)** Bout duration frequencies for a stimlus strength of 94.35Hz are plotted on a log-scale with an exponential fit shown (orange). **B)** Same as (A) but for a stimulus of strength 377.4Hz. **C)** Same as (A,B) but for a stimulus of strength 660.45Hz.

**Supplementary Figure 4.**
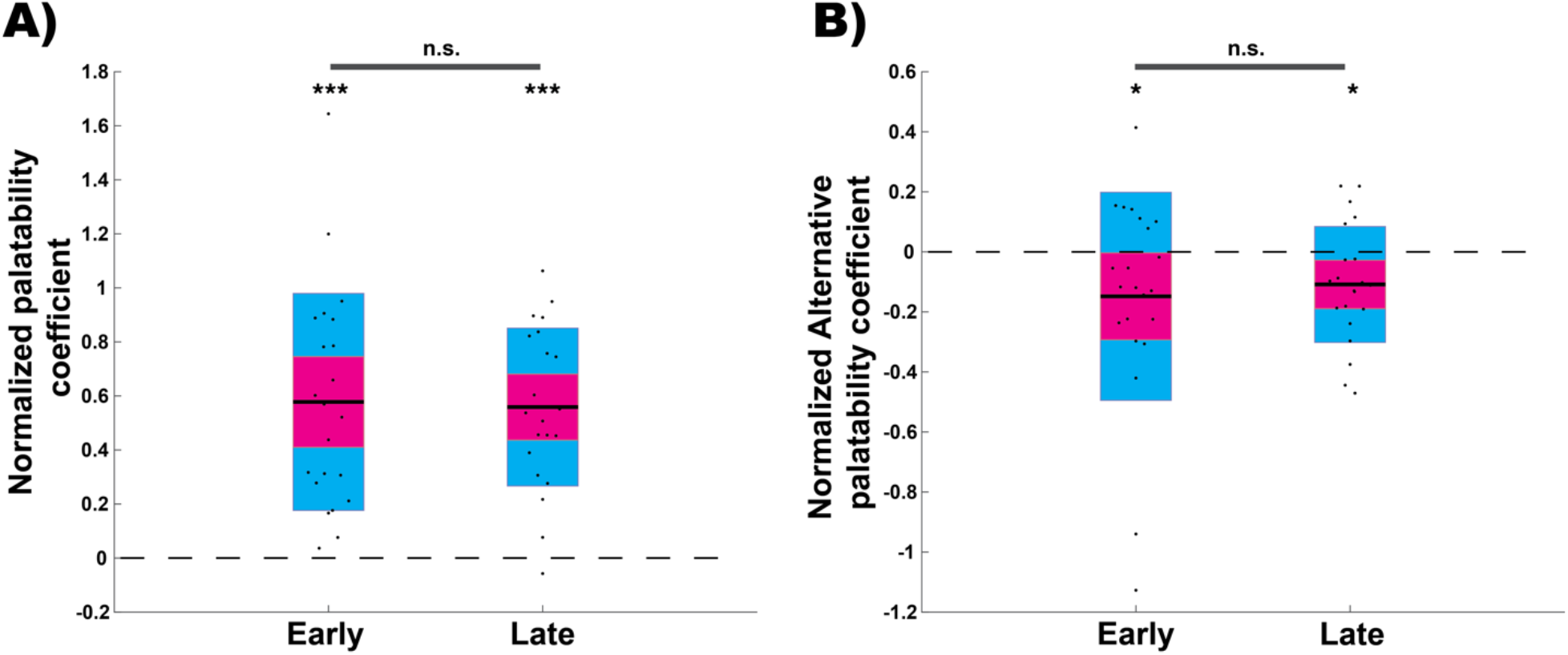
Normalized regression coefficients for early and late bouts defined using the 200ms ILI criterion. **A)** Normalized regression coefficients for current solution palatability for bouts in the early or late portion of the session. Current palatability coefficients were significantly positive for both the early (right-tailed: z = 4.09, p < .001) and late (right-tailed: z = 4.06, p < .001) portions of the task. Coefficients were not significantly different across portions of the session (paired: z = .016, p = .987). **B)** Same as (A) but for the alternative solution’s palatability. Normalized coefficients were significantly negative for both early (left-tailed: z = −1.85, p = .032) and late portions of the session (left-tailed: z = −2.2, p = .014). Coefficients were not significantly different across portions of the session (paired: z = −.11, p = .91).

**Supplementary Figure 5.**
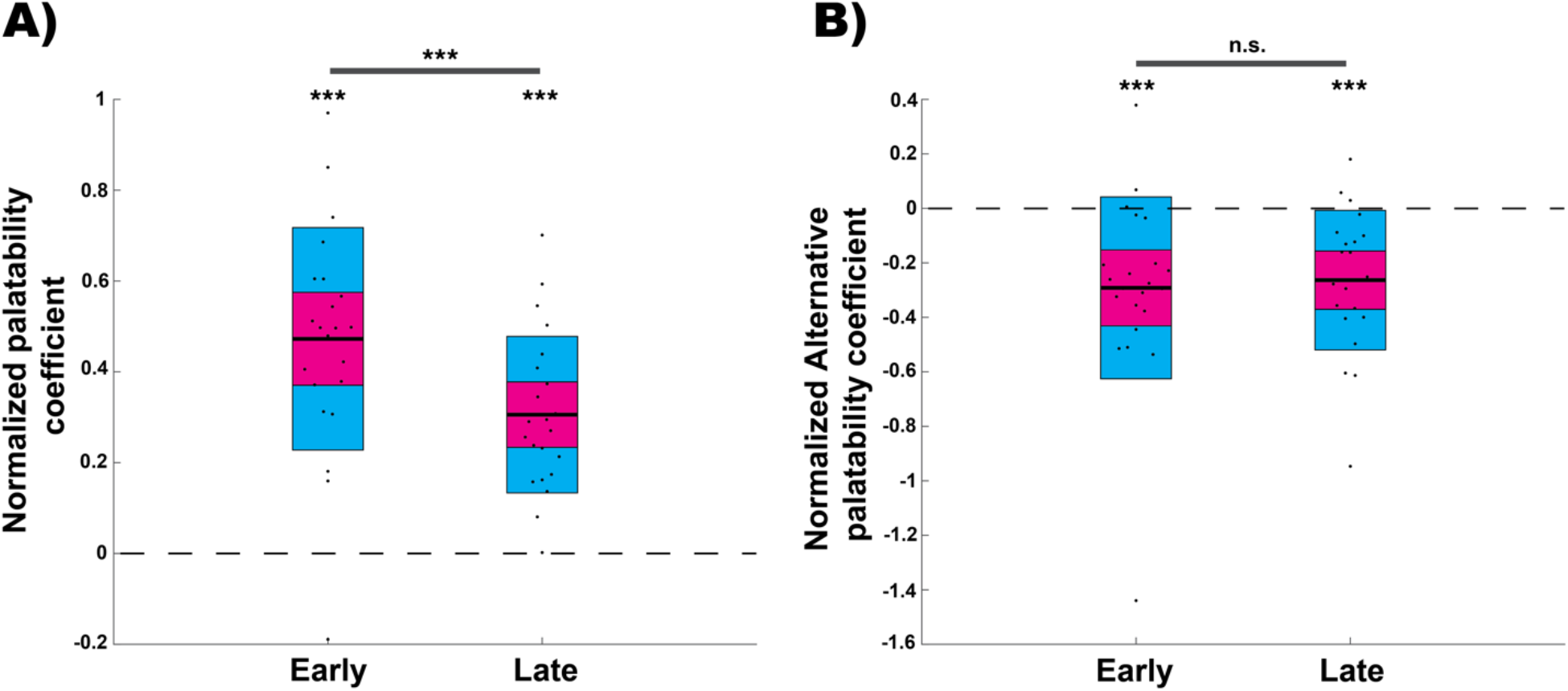
Normalized current/alternative palatability regression coefficients for early/late bouts when using the altered rank-order of palatability (Pal(0.01M) = 1.3xPal(0.1M)). **A)** Coefficients during the early portion (right-tailed Wilcoxon signed-rank: z = 3.99, p < .001) and late portion (right-tailed Wilcoxon signed-rank: z = 4.09, p < .001) were significantly positive. Normalized palatability coefficients for the early part of the session were significantly more positive than those for the late part (right-tailed Wilcoxon signed-rank: z = 3.18, p < .001). **B)** Normalized alternative palatability coefficients during the early portion (left-tailed Wilcoxon signed-rank: z = −3.38, p < .001) and late portion (left-tailed Wilcoxon signed-rank: z = −3.6, p < .001) of the task were significantly negative. There is no difference between coefficients in the early and late portions of the session (paired two-tailed Wilcoxon signed-rank: z = −.5, p = .61).

